# Lifespan and fecundity impacts of reduced insulin signalling can be directed by mito-nuclear epistasis in *Drosophila*

**DOI:** 10.1101/2025.09.04.674084

**Authors:** Rita Ibrahim, Christin Froschauer, Susanne Broschk, David R Sannino, Adam J Dobson

## Abstract

The changing demography of human populations has motivated a search for interventions that promote healthy ageing, and especially for evolutionarily-conserved mechanisms that can be studied in lab systems to generate hypotheses about function in humans. Reduced Insulin/IGF signalling (IIS) is leading example, which can extend healthy lifespan in a range of animals; but whether benefits and costs of reduced IIS vary genetically within species is under-studied. This information is critical for any putative translation. Here, in *Drosophila*, we test for genetic variation in lifespan response to a dominant-negative form of the insulin receptor, along with a metric of fecundity to evaluate corollary fitness costs/benefits. We also partition genetic variation between DNA variants in the nucleus (nDNA) and mitochondrial DNA (mtDNA), in a fully-factorial design that allows us to assess “mito-nuclear” epistasis. We show that reduced IIS can have either beneficial or detrimental effects on lifespan, depending on the combination of mtDNA and nDNA. This suggests that, while insulin signalling has a conserved effect on ageing among species, intraspecific effects can vary genetically, and the combination of mtDNA and nDNA can act as gatekeeper.

## Main text

Human life expectancy has increased over recent centuries, increasing the prevalence of age-related diseases such as cardiovascular disease, neurodegeneration, and cancer (Singh et al. 2019; Niccoli & Partridge 2012; Li et al. 2021). This has motivated a search for fundamental mechanisms of ageing. The insulin/IGF signalling (IIS) pathway is a key regulator of ageing, from invertebrates to mammals (Ziv & Hu 2011; Kenyon et al. 1993; Clancy et al. 2001; Selman et al. 2008; Ikeya et al. 2009; Broughton et al. 2005; Dorman et al. 1995), and naturally-occurring variants in this pathway are associated with human longevity (Singh et al. 2019; Chung et al. 2010). This evolutionarily-conserved role has generated considerable interest in better understanding mechanistically how and why IIS impacts ageing. However, IIS activity has adaptive functions early in life, e.g. in transcription, translation, growth, metabolic regulation, and mitogenesis (Li et al. 2014), and so any putative anti-ageing intervention impairing the pathway must balance potential benefits against the probable costs to these traits. This necessitates understanding how costs and benefits vary, and what underpins this variation.

Ageing is subject to standing genetic variation (Yuan et al. 2020; Vaught et al. 2020; Durham et al. 2014), and consequently we expect individual variation in responses to anti-ageing interventions. However, studies documenting such variation are scarce. We have shown genetic variation in *Drosophila* lifespan and mortality response to rapamycin (Ibrahim et al. 2024), and in multiple species variation in effects of dietary restriction has also been well documented (McCracken et al. 2020; Gautrey & Simons 2022; Durham et al. 2014; Green et al. 2022; Liao et al. 2013; Rikke et al. 2010; Francesco et al. 2024). With reference to IIS, mouse strains vary in their lifespan response to IGF-1R mutation (Xu et al. 2014; Selman & Swindell 2018; Bokov et al. 2011), but otherwise we are not aware of systematic investigations, nor of candidate mechanisms. Characterising and understanding such variation is key to unlocking any putative translational potential of IIS impairment.

Accumulating evidence suggests that a significant amount of genetic variation is not caused by additive effects of variants, but rather by epistatic interactions between independent loci (Boyle et al. 2017). mtDNA and nDNA are inherited partly independently, because mtDNA is only inherited maternally, while nDNA is inherited from both parents: this generates “mito-nuclear” epistasis for diverse traits including lifespan, reproductive fitness, development, stress tolerance and metabolism (Garlovsky et al. 2025; Dowling & Wolff 2023). Here, we ask whether mito-nuclear epistasis modulates response to IIS in female *Drosophila*, testing whether impacts on lifespan are universally positive or variable.

We impaired insulin signalling in *Drosophila* by ubiquitous expression of a dominant-negative form of the insulin receptor *InR (InR*^*DN*^), which has a well-established lifespan-extending effect, in studies performed predominantly in the *Dahomey* background (Ikeya et al. 2009; Slack et al. 2011; Dobson et al. 2019; Bolukbasi et al. 2017). We drove *InR*^*DN*^ expression ubiquitously using the *Daughterless-GeneSwitch (DaGS)* driver, activated by feeding on a chemical inducer, RU486. We established a new panel of fly lines to evaluate how the lifespan output of *InR*^*DN*^ varies genetically, and how it can be shaped by variation in mtDNA, nDNA and mito-nuclear epistasis. We refer to the various combinations of mtDNA and nDNA as mitonucleogenotypes (Dobson et al. 2023; Ibrahim et al. 2024). We confirmed that the stocks we used to found this panel were free of the cytoplasmic endosymbiont *Wolbachia*, which could confound our experiment by coinheritance with mtDNA (Figure S1A). We took flies from the *Canton-S, Dahomey* and *w1118* backgrounds (subsequently C, D and E, respectively), and replicated a design that we and others have used previously to generate mitonuclear variation (Figure S1B) (Garlovsky et al. 2025; Vaught et al. 2020; Dobson et al. 2023; Ibrahim et al. 2024), introgressing nDNAs into mtDNA backgrounds by crossing males (nDNA donor) to females with the desired mtDNA. We iterated this procedure six times, which should eliminate >95% of residual nDNA variation from F0 mothers. We used a fully-factorial design, generating all nine possible combinations of mtDNA and nDNA (i.e. nine mitonucleogenotypes) in triplicated parallel backcrosses (i.e. 27 lines), into which we then crossed *DaGS* and *UAS-InR*^*DN*^ separately (i.e. generating 54 lines), from donors with matched nDNA (Figure S1B). We homozygosed the offspring and then crossed *DaGS* and *UAS-InR*^*DN*^ flies from the same lines, to finally nine mitonucleogenotypes, each triplicated to generate 27 lines altogether, all of which were heterozygous for each transgene and could therefore inducibly express *InR*^*DN*^ ubiquitously. We fed three day-old adult females RU486 (henceforth RU), to induce ubiquitous *InR*^*DN*^ expression, and we scored subsequent survival, and egg laying after one week of induction.

We compared survival in the presence/absence of RU for each mitonucleogenotype, (Figure 1A), and analysed the data using parametric survival models (PSM). Overall, a three-way interaction of RU, mtDNA and nDNA was apparent (LR ChiSq=25.41, Df=4, p<0.0001). To identify what drove this interaction, we fit a second model which revealed mitonucleogenotype-specific effects of RU (mitonucleogenotype:RU LR ChiSq=101.54, Df=8, p<2.2e-16). We used post-hoc tests (estimated marginal means, henceforth EMmeans) to identify specifically how different combinations of mtDNA and nDNA led to distinct survival outcomes, both for the pool of three replicate populations (Figure 1A, Figure S1B) and for each of the three constituent replicates mitonucleogenotype. We accepted the overall result in the analyses of the three pooled replicates only if the majority of replicate lines (i.e. two of three) were in agreement. This approach gives a false discovery rate of 11%, i.e. the square of the probability (1/3) of randomly detecting one of the three possible results (i.e. lifespan increased, lifespan decreased, no effect), squared (1/3 ^*^ 1/3 = 0.11). This conservative methodology led us to reject a result from analysing the pool of *CE* populations (Figure S2A), because we were only able to detect an effect in population *CE3* (Figure S2B). Nevertheless, the overall result of the pooled analysis – a tripartite interaction between RU, mtDNA and nDNA – was robust to this filter. Specifically, the impact of RU was modulated by nDNA in the presence of mtDNAs *C* or *E*, shortening lifespan with nDNA *C* but extending lifespan with nDNA *D*; but mtDNA *D* blocked any response to RU regardless of nDNA (Figure 1A). Thus, the results indicated that (A) lifespan response to reduced insulin signalling can vary genetically, and (B) mito-nuclear epistasis can play a role.

**Figure 1.**
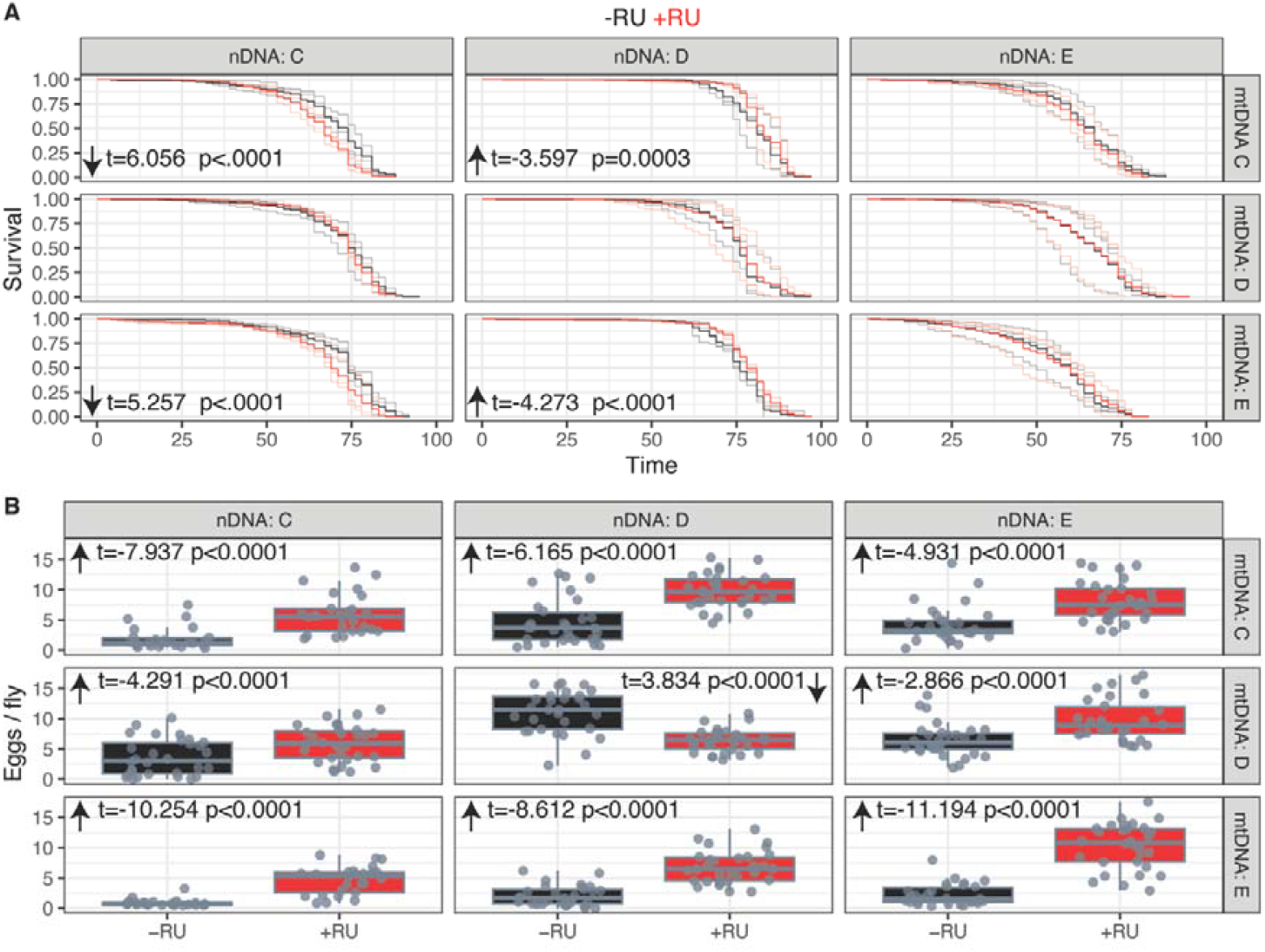
Mito-nuclear epistasis directs the effect of reduced insulin signalling on lifespan and egg laying. Arrows on all facets indicate effects of RU relative to control. **A**. Kaplan-Meier survival plots of flies DaGS/UAS-InR^DN^ flies, faceted by mitonucleogenotypes. Survival of control flies shown in black, survival of flies fed 200µM RU486 shown in red. Thin lines give survival of replicate populations, thicker lines give grand mean across replicates. Parametric survival model (PSM), mitonucleogenotype:RU LR ChiSq=101.54, Df=8, p<2.2e-16. Statistics given on panels when significant effects of RU were detected in post-hoc (EMMeans) analysis of PSM, negative t-ratio indicates longer lifespan in RU-fed flies than controls. **B**. Boxplots show eggs laid per DaGS > UAS-InR^DN^ female (nine days old), in the presence (red) or absence (black) of RU486. The boxplots represent the median, first and third quartiles and 5^th^ and 95^th^ percentiles. Statistics given on panels from post-hoc tests on a negative binomial mixed model (mitonucleogenotype:RU Chisq=180.601, df=8, p<2.2e-16).

Fecundity was decreased by feeding RU to flies bearing mtDNA *D* and nDNA *D*. This result was consistent with the theoretical expectation that lifespan should negatively correlate fitness. However, contrary to expectation, in every other mitonucleogenotype under study, fecundity was increased by RU (Figure 1B). These differences manifested as a mitonucleogenotype-specific effect of RU (Negative binomial GLMM, mitonucleogenotype:RU Chisq=180.601, df=8, p<2.2e-16), and a significant RU:mtDNA:nDNA interaction (Negative Binomial GLM, ChiSq=14.29, df=4, p<0.01). Together with the preceding lifespan data, the fecundity results suggest that fitness costs of *InR*^*DN*^ expression are not obligate, and that in fact the relationship between lifespan cost and fitness benefit can be shaped by mito-nuclear epistasis.

Impaired IIS clearly has capacity to promote healthy ageing, with effects reported widely across species (Partridge et al. 2010; Selman et al. 2008; Clancy et al. 2001; Kenyon 2010; Zhang et al. 2013; Murphy et al. 2003). It seems unlikely that beneficial effects could have been reported in species separated by hundreds of millions of years of evolutionary divergence if fundamental processes were not at play. Nevertheless, our present results, within just one species, suggest that one means of reducing IIS can have effects that vary within species. These results also complement our previous results, showing that the magnitude and timing of the effect of a pharmacological anti-ageing intervention (rapamycin)(Ibrahim et al. 2024). Together, these studies imply that benefits of interventions with conserved effects are not necessarily universal, and that their benefits may be conditional. We manipulated genotype (specifically mitonucleogenotype), showing that different genotypes raised under standardised conditions exhibited different responses, but one intriguing possibility is that the capacity to benefit from reduced IIS may be subject to interactions between genotype and other variables, e.g. environment, diet composition, timing of induction. It is possible that many genotypes have inherent capacity to benefit from anti-ageing interventions, but context and conditions must be tailored appropriately. Such interactions could allow genotypes that experienced neutral or deleterious effects in the present study to show benefits in other contexts. Thus, the results imply that better understanding the context-specificity of responses to anti-ageing interventions should be a priority for the field. The basis for variation in response is likely complex, potentially incorporating interactions among genetics, nutrition, environment etc; which will be difficult to disentangle. We therefore suggest that research should focus on characterising the emergent properties that predispose an individual to benefit from a given intervention.

We have shown specifically that the impact of impaired IIS is influenced by the “lock and key” of mito-nuclear epistasis. Emerging questions now are why should mtDNA, nDNA, and their epistatic interaction impact response to reduced IIS? Is this because IIS alters mitochondrial function in ways that depend on nDNA variants? Or do mito-nuclear effects alter how IIS impacts cellular function? Mechanistic work is now required to address these issues. Furthermore, the present results complement and extend our previous finding that anti-ageing impacts of rapamycin were more universally beneficial, but nevertheless shaped by mito-nuclear epistasis. We therefore think it likely that other anti-ageing interventions will be subject to genetic variation, and likely mito-nuclear variation.

Altogether, our results provide new evidence that interventions that modulate ageing among species can nevertheless be subject to genetic variation within species, which can be underpinned by mtDNA variation, nDNA variation, and epistatic mito-nuclear variation. Importantly, the intervention used here can lead to directionally different outcomes, implying that (1) we do not understand genetic variation contextualises signalling, and (2) long-term translational goals should include precisely matching interventions to those most likely to benefit.

## Materials and Methods

Flies bearing the *w1118* mutation in the *CantonS, “w1118”* and *OregonR* backgrounds were gifts from the lab of Julian Dow, University of Glasgow. Flies bearing the *w1118* mutation in the *Dahomey* background were gifts from the UCL Institute of Healthy Ageing. Flies bearing *DaGS* and *UAS-InR*^*DN*^ transgenes were gifts from Cathy Slack, University of Warwick. A panel of 9 mitonucleogenotypes established as previously (Ibrahim et al. 2024; Dobson et al. 2023; Vaught et al. 2020) with 6 rounds of backcrossing, as described in Figure S1. Triplicating each mitonucleogenotype generated 27 populations altogether. Transgenes were backcrossed into the three respective nDNA backgrounds, crossed to mitonucleotypes bearing the same nDNA background, and homozygosed. Virgin females bearing *DaGS* (n=45) from each of the 27 populations were crossed for 48h in egg laying cages to males from the same populations bearing *UAS-InR*^*DN*^. Eggs were collected after an overnight egg lay, suspended in PBS, and 20µl suspension was pipetted onto 60ml sugar-yeast-agar (1xSYA) medium containing 10% (w/v) brewer’s yeast (MP Biomedicals SR03010, LOT No. S4707), 1.5% (w/v) agar (Sigma A7002), 5% (w/v) sucrose (Fisher BP220-10), 0.3% nipagin (10% in EtOH), 0.3% propionic acid (Bass et al. 2007). Upon eclosion as adults, flies were transferred to fresh food and allowed to mate 48h before males were discarded. Females were fed food containing 200µM RU486 (Cayman Chemical CAY10006317) to induce *InR*^*DN*^ expression, with controls fed EtOH as a vehicle control. Survival was scored thrice weekly until all flies had died. Overnight egg laying was assayed after one week feeding on RU486.

Data were analysed in R 4.4.2. Survival data were analysed with a parametric survival model (rms::PSM). A logistic distribution was determined to best fit the data by comparing AIC values. ANOVA tables were calculated with car::Anova, using Type-3 tests. Post-hoc tests were calculated with the EMmeans library.

## Author Contributions

RI, CF, SB and DRS acquired data. RI and AJD performed data analysis. AJD conceived and supervised the study, and wrote the manuscript.

## Acknowledgments

This work was supported by a University of Glasgow Lord Kelvin Adam Smith Scholarship to RI, a UKRI Future Leaders Fellowship (MR/S033939/1 and MR/Y019660/1) to AJD, and a Lord Kelvin Adam Smith Fellowship to AJD. We thank Nathan Woodling and Colin Selman for comments which improved the manuscript; and Klaus Reinhardt for ongoing support and constructive discussion.

## Conflict of interest statement

The authors declare no conflicts of interest.

## Data availability statement

The data that support the findings of this study are openly available at https://github.com/dobdobby/InR-mito-nuclear

---

**Figure S1.**
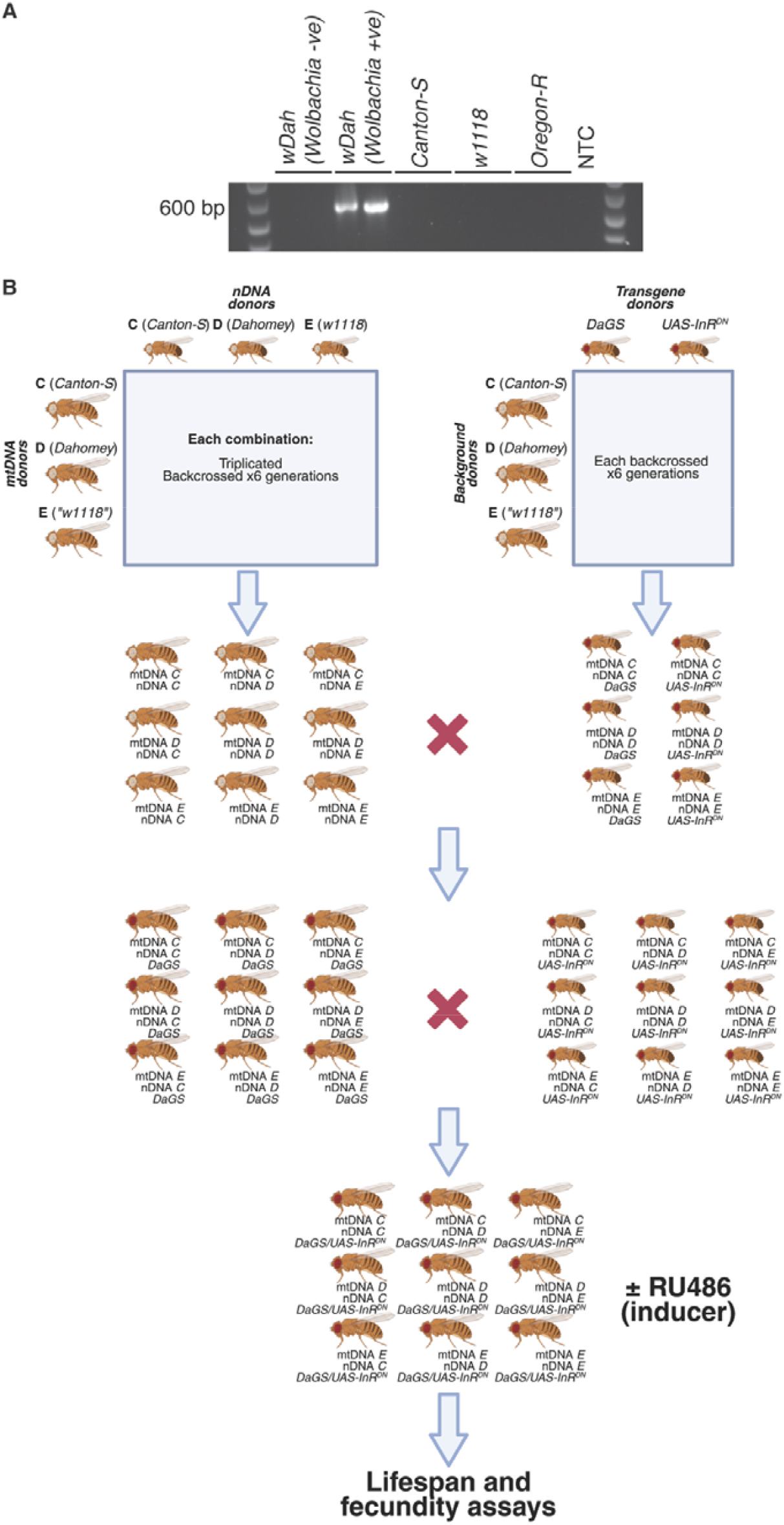
Establishing new *Drosophila* populations to enable ubiquitous expression of dominant-negative *InR* in varied mitonucleogenotypes. **A**. Lab populations were tested for presence of *Wolbachia* endosymbionts, by PCR using species-specific 16s primers. *Dahomey* flies known to be *Wolbachia* positive or negative (Ikeya et al. 2009) were included as positive and negative controls, respectively. *Canton-S, w1118* and *Oregon-R* populations in our lab were also found to be *Wolbachia*-negative. NTC=no-template control. **B**. 27 newly-established populations show mito-nuclear variation in egg laying. Populations were derived from *Canton-S (C), Wolbachia*-negative *Dahomey (D)*, and *w1118 (E)*, by backcrossing nDNA over mtDNA (see panel C) in a fully-factorial design, generating nine mitonucleogenotypes, each in three replicate backcrosses. For each population the first letter represents mtDNA, second represents nDNA, and 1-3 represents independent replicate backcrosses (noting that 1-3 are independent across the nine mitonucleogenotypes), e.g. *CD1* = one replicate of mtDNA *C*, nDNA *D*. **C**. Combining functional genetics with mitonuclear variation. Nine mitonucleogenotypes were established (as shown in panels A and B). In parallel, *DaGS* and *UAS-InR*^*DN*^ transgenes were backcrossed into corresponding ancestral *C, D* and *E* backgrounds. Transgene-bearing flies were then crossed to nDNA*-* matched mitonuclear flies (e.g. transgenes in nDNA *C* background were crossed to lines *CC1-3, DC1-3, EC1-3*; transgenes in nDNA *E* background were crossed to lines *CE1-3, DE1-3, EE1-3*, etc). Resulting heterozygotes were homozygosed, generating flies from nine mitonucleogenotypes bearing *DaGS* or *UAS-InR*^*DN*^, each in triplicate. Flies were crossed to finally generate nine mitonucleogenotypes (each triplicated) bearing *DaGS/UAS-InR*^*DN*^.

**Figure S2.**
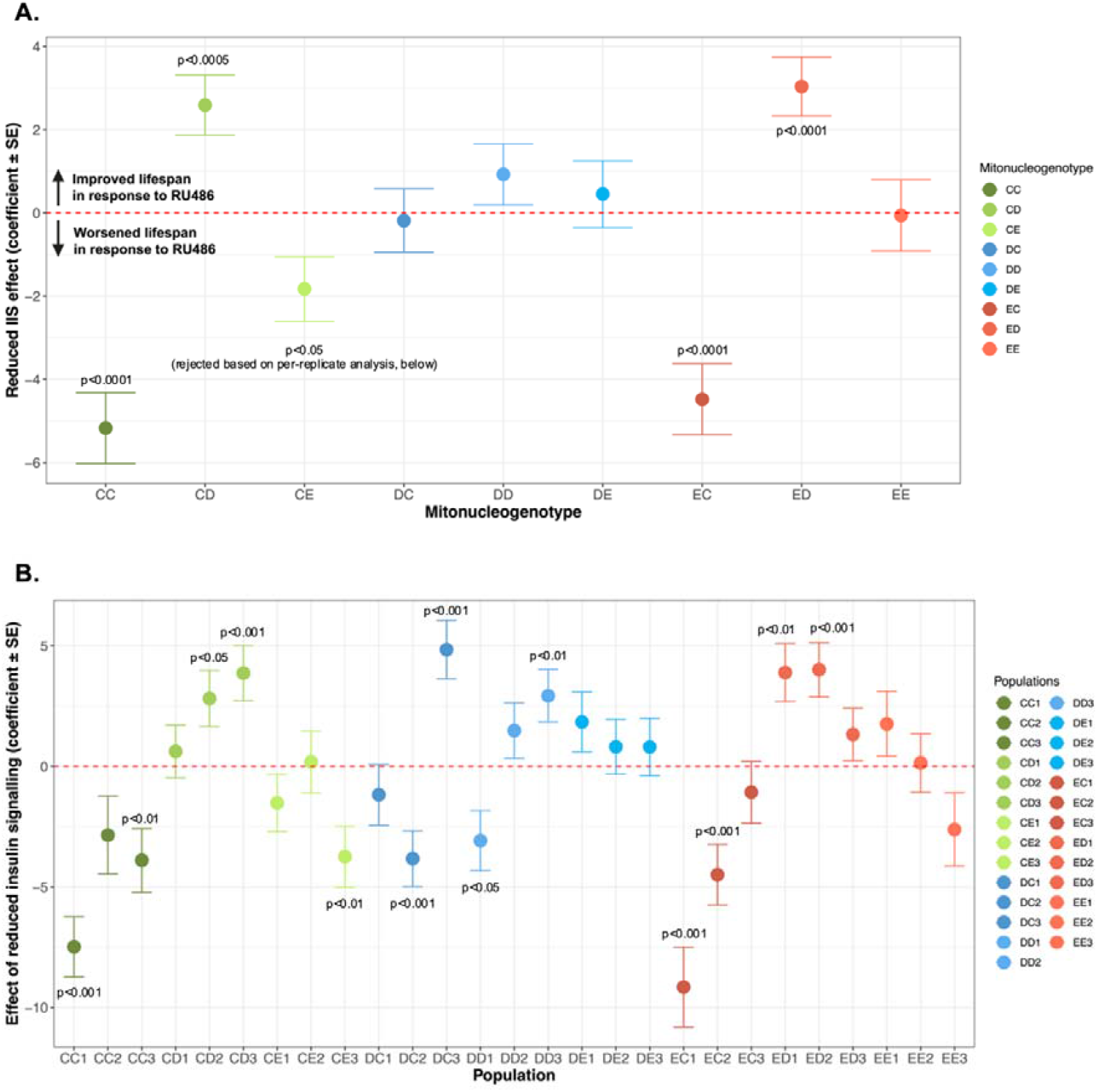
Lifespan effects of *InR*^*DN*^ in nine mitonucleogenotypes and 27 constituent replicates. **A**. Pooled populations - Coefficients from post-hoc analysis (impact of RU, stratified by replicate population), of EMMeans ±SE of a PSM (logistic distribution). **B**. Constituent replicates of the pooled populations - Coefficients from post-hoc analysis (impact of RU, stratified by mtDNA and nDNA), of EMMeans ± SE. Coefficients reflect the impact of *InR*^*DN*^ induction on lifespan. Coefficients are multiplied -1^*^x, such that higher values correspond to extended lifespan. Red horizontal represents no impact of *InR*^*DN*^.

## Notes

### Competing Interest Statement

The authors have declared no competing interest.

## Bibliography

Bass TM, Grandison RC, Wong R, Martinez P, Partridge L & Piper MDW (2007) Optimization of Dietary Restriction Protocols in Drosophila. Journals Gerontology Ser 62, 1071–1081.

Bokov AF, Garg N, Ikeno Y, Thakur S, Musi N, DeFronzo RA, Zhang N, Erickson RC, Gelfond J, Hubbard GB, Adamo ML & Richardson A (2011) Does Reduced IGF-1R Signaling in Igf1r +/− Mice Alter Aging? PLoS ONE 6, e26891.

Bolukbasi E, Khericha M, Regan JC, Ivanov DK, Adcott J, Dyson MC, Nespital T, Thornton JM, Alic N & Partridge L (2017) Intestinal Fork Head Regulates Nutrient Absorption and Promotes Longevity. Cell Reports 21.

Boyle EA, Li YI & Pritchard JK (2017) An Expanded View of Complex Traits: From Polygenic to Omnigenic. Cell 169, 1177–1186.

Broughton SJ, Piper MD, Ikeya T, Bass TM, Jacobson J, Driege Y, Martinez P, Hafen E, Withers DJ, Leevers SJ & Partridge L (2005) Longer lifespan, altered metabolism, and stress resistance in Drosophila from ablation of cells making insulin-like ligands. Proceedings of the National Academy of Sciences of the United States of America 102, 3105–3110.

Chung W-H, Dao R-L, Chen L-K & Hung S-I (2010) The role of genetic variants in human longevity. Ageing Res. Rev. 9, S67–S78.

Clancy DJ, Gems D, Harshman LG, Oldham S, Stocker H, Hafen E, Leevers SJ & Partridge L (2001) Extension of Life-Span by Loss of CHICO, a Drosophila Insulin Receptor Substrate Protein. Science 292, 104–106.

Dobson AJ, Boulton-McDonald R, Houchou L, Svermova T, Ren Z, Subrini J, Vazquez-Prada M, Hoti M, Rodriguez-Lopez M, Ibrahim R, Gregoriou A, Gkantiragas A, Bähler J, Ezcurra M & Alic N (2019) Longevity is determined by ETS transcription factors in multiple tissues and diverse species. PLOS Genetics 15, e1008212.

Dobson AJ, Voigt S, Kumpitsch L, Langer L, Voigt E, Ibrahim R, Dowling DK & Reinhardt K (2023) Mitonuclear interactions shape both direct and parental effects of diet on fitness and involve a SNP in mitoribosomal 16s rRNA. PLOS Biol. 21, e3002218.

Dorman JB, Albinder B, Shroyer T & Kenyon C (1995) The age-1 and daf-2 genes function in a common pathway to control the lifespan of Caenorhabditis elegans. Genetics 141, 1399–1406.

Dowling DK & Wolff JN (2023) Evolutionary genetics of the mitochondrial genome: insights from Drosophila. GENETICS 224, iyad036.

Durham MF, Magwire MM, Stone EA & Leips J (2014) Genome-wide analysis in Drosophila reveals age-specific effects of SNPs on fitness traits. Nature Communications 5.

Francesco AD, Deighan AG, Litichevskiy L, Chen Z, Luciano A, Robinson L, Garland G, Donato H, Vincent M, Schott W, Wright KM, Raj A, Prateek GV, Mullis M, Hill WG, Zeidel ML, Peters LL, Harding F, Botstein D, Korstanje R, Thaiss CA, Freund A & Churchill GA (2024) Dietary restriction impacts health and lifespan of genetically diverse mice. Nature 634, 684–692.

Garlovsky MD, Dobler R, Guo R, Voigt S, Dowling DK & Reinhardt K (2025) Testing for age- and sex-specific mitonuclear epistasis in Drosophila. Evolution, qpaf096.

Gautrey SL & Simons MJP (2022) Amino Acid Availability Is Not Essential for Life-Span Extension by Dietary Restriction in the Fly. J. Gerontol.: Ser. A 77, 2181–2185.

Green CL, Lamming DW & Fontana L (2022) Molecular mechanisms of dietary restriction promoting health and longevity. Nat. Rev. Mol. Cell Biol. 23, 56–73.

Ibrahim R, Martinez MB & Dobson AJ (2024) Rapamycin’s lifespan effect is modulated by mito□nuclear epistasis in Drosophila. Aging Cell 23, e14328.

Ikeya T, Broughton S, Alic N, Grandison R & Partridge L (2009) The endosymbiont Wolbachia increases insulin/IGF-like signalling in Drosophila. Proceedings of the Royal Society B: Biological Sciences 276, 3799–3807.

Kenyon C (2010) A pathway that links reproductive status to lifespan in Caenorhabditis elegans. Annals of the New York Academy of Sciences 1204, 156–162.

Kenyon C, Chang J, Gensch E, Rudner A & Tabtiang R (1993) A C. elegans mutant that lives twice as long as wild type. Nature 366, 461.

Li CR, Guo D & Pick L (2014) Independent signaling by Drosophila insulin receptor for axon guidance and growth. Front. Physiol. 4, 385.

Li Z, Zhang Z, Ren Y, Wang Y, Fang J, Yue H, Ma S & Guan F (2021) Aging and age□related diseases: from mechanisms to therapeutic strategies. Biogerontology 22, 165–187.

Liao C-Y, Johnson TE & Nelson JF (2013) Genetic variation in responses to dietary restriction — An unbiased tool for hypothesis testing. Exp. Gerontol. 48, 1025–1029.

McCracken AW, Adams G, Hartshorne L, Tatar M & Simons MJP (2020) The hidden costs of dietary restriction: Implications for its evolutionary and mechanistic origins. Sci Adv 6, eaay3047.

Murphy CT, McCarroll SA, Bargmann CI, Fraser A, Kamath RS, Ahringer J, Li H & Kenyon C (2003) Genes that act downstream of DAF-16 to influence the lifespan of Caenorhabditis elegans. Nature 424.

Niccoli T & Partridge L (2012) Ageing as a Risk Factor for Disease. Current Biology 22.

Partridge L, Alic N, Bjedov I & Piper M (2010) Ageing in Drosophila: The role of the insulin/Igf and TOR signalling network. Experimental Gerontology 46.

Rikke BA, Liao C-Y, McQueen MB, Nelson JF & Johnson TE (2010) Genetic dissection of dietary restriction in mice supports the metabolic efficiency model of life extension. Exp Gerontol 45, 691–701.

Selman C, Lingard S, Choudhury AI, Batterham RL, Claret M, Clements M, Ramadani F, Okkenhaug K, Schuster E, Blanc E, Piper MD, Al□Qassab H, Speakman JR, Carmignac D, Robinson ICA, Thornton JM, Gems D, Partridge L & Withers DJ (2008) Evidence for lifespan extension and delayed age–related biomarkers in insulin receptor substrate 1 null mice. Faseb J 22, 807–818.

Selman C & Swindell WR (2018) Putting a strain on diversity. EMBO J. 37, EMBJ2018100862.

Singh PP, Demmitt BA, Nath RD & Brunet A (2019) The Genetics of Aging: A Vertebrate Perspective. Cell 177, 200–220.

Slack C, Giannakou ME, Foley A, Goss M & Partridge L (2011) dFOXO□independent effects of reduced insulin□like signaling in Drosophila. Aging Cell 10, 735–748.

Vaught RC, Voigt S, Dobler R, Clancy DJ, Reinhardt K & Dowling DK (2020) Interactions between cytoplasmic and nuclear genomes confer sex□specific effects on lifespan in Drosophila melanogaster. J Evolution Biol 33, 694–713.

Xu J, Gontier G, Chaker Z, Lacube P, Dupont J & Holzenberger M (2014) Longevity effect of IGF□1R+/− mutation depends on genetic background□specific receptor activation. Aging Cell 13, 19–28.

Yuan R, Musters CJM, Zhu Y, Evans TR, Sun Y, Chesler EJ, Peters LL, Harrison DE & Bartke A (2020) Genetic differences and longevity□related phenotypes influence lifespan and lifespan variation in a sex□specific manner in mice. Aging Cell 19, e13263.

Zhang P, Judy M, Lee S-J & Kenyon C (2013) Direct and Indirect Gene Regulation by a Life-Extending FOXO Protein in C. elegans: Roles for GATA Factors and Lipid Gene Regulators. Cell Metabolism 17.

Ziv E & Hu D (2011) Genetic variation in insulin/IGF-1 signaling pathways and longevity. Ageing Res. Rev. 10, 201–204.

